# Swine H1N1 influenza virus variants with enhanced polymerase activity and HA stability promote airborne transmission in ferrets

**DOI:** 10.1101/2021.09.22.461336

**Authors:** Meng Hu, Jeremy C. Jones, Balaji Banoth, Chet Raj Ojha, Jeri Carol Crumpton, Lisa Kercher, Robert G. Webster, Richard J. Webby, Charles J. Russell

## Abstract

Understanding how animal influenza A viruses (IAVs) acquire airborne transmissibility in humans and ferrets is needed to prepare for and respond to pandemics. Here, we investigated in ferrets the replication and transmission of swine H1N1 isolates P4 and G15, whose majority population had decreased polymerase activity and poor HA stability, respectively. For both isolates, a minor variant was selected and transmitted in ferrets. Polymerase-enhancing variant PA-S321 airborne-transmitted and propagated in one ferret. HA-stabilizing variant HA1-S210 was selected in all G15-inoculated ferrets and was transmitted by contact and airborne routes. With an efficient polymerase and a stable HA, the purified minor variant G15-HA1-S210 had earlier and higher peak titers in inoculated ferrets and was recovered at a higher frequency after airborne transmission than P4 and G15. Overall, HA stabilization played a more prominent role than polymerase enhancement in the replication and transmission of these viruses in ferrets. The results suggest pandemic risk-assessment studies may benefit from deep sequencing to identify minor variants with human-adapted traits.

**IMPORTANCE:** Diverse IAVs circulate in animals, yet few acquire the viral traits needed to start a human pandemic. A stabilized HA and mammalian-adapted polymerase have been shown to promote the adaptation of IAVs to humans and ferrets (the gold-standard model for IAV replication, pathogenicity, and transmissibility). Here, we used swine IAV isolates of the gamma lineage a model to investigate the importance of HA stability and polymerase activity in promoting replication and transmission in ferrets. These are emerging viruses that bind to both *α*-2,6- and *α*-2,3-linked receptors. Using isolates containing mixed populations, a stabilized HA was selected within days in inoculated ferrets. An enhanced polymerase was also selected and propagated after airborne transmission to a ferret. Thus, HA stabilization was a stricter requirement, yet both traits promoted transmissibility. Knowing the viral traits needed for pandemic potential, and the relative importance of each, will help identify emerging viruses of greatest concern.

## INTRODUCTION

IAVs circulate in numerous species, and it remains challenging to identify animal-origin IAVs at greatest risk to cause a pandemic in humans. In addition to bats (1–3), IAVs originate from a reservoir of wild aquatic birds (4). From wild birds, avian IAVs can infect and either directly or indirectly become endemic in wild and domestic birds, swine, humans, and other domestic or aquatic mammals (5, 6). IAVs have eight RNA gene segments that encode for over a dozen proteins including an RNA-dependent RNA polymerase (RdRp) complex (PB1, PB2, and PA) and the surface glycoproteins hemagglutinin (HA) and neuraminidase (NA) (7, 8). Antigenically diverse HA and NA subtypes have been identified and currently number from H1-H18 and N1-N11. H1N1 caused the 1918 Spanish influenza pandemic, circulated in humans until 1957, became endemic in swine, reemerged in humans in 1977, and caused the 2009 pandemic (pH1N1) (8, 9). In 1957, H1N1 was supplanted in humans by the H2N2 Asian influenza pandemic. H2N2 was supplanted by the H3N2 Hong Kong influenza pandemic in 1968. H3N2, pH1N1, and influenza B viruses cause seasonal influenza in humans. H1N1, H1N2, and H3N2 viruses are currently endemic in swine (10–12).

H1 swine viruses have been divided into six clades: alpha (1A.1), beta (1A.2), gamma (1A.3.3.3), H1pandemic (1A.3.3.2), delta1 (1B.2.2), and delta2 (1B.2.1) (13, 14). Swine gamma and pandemic clades diverged approximately twenty years ago and have caused numerous human infections (15). Currently, H1 gamma clade is endemic in swine and frequently reassorts with human pandemic H1N1 isolates (16), which may result in increased transmissibility in swine or humans. Preparing for and responding to future human pandemics requires an understanding of viral traits and host markers associated with human adaptation so that high-risk emerging viruses can be identified, and countermeasures can be taken.

IAV adaptations that have been associated with human-to-human transmission include HA receptor-binding specificity, HA stability, polymerase complex efficiency, HA-NA balance, NA stalk length, interferon antagonism, and filamentous virus morphology (5, 17). During viral entry, the HA protein binds sialic acid-containing receptors. This triggers virion endocytosis and acidification, which induces irreversible HA structural changes that cause membrane fusion. Human-adapted IAVs contain HA proteins that preferentially bind *α*-2,6-linked sialic acid (SA) receptors over *α*-2,3-linked ones, which are preferred by avian IAVs (8, 18–22). In human- and ferret-adapted IAVs, the avidity of the HA protein for cell-surface glycan receptors is balanced with the substrate selectivity and catalytic rate of the NA protein (17, 23–26). Decreased HA activation pH in the context of H5N1 viruses reduces replication and transmission in avian species but increases nasal replication and is required for airborne transmission in ferrets (27–34). Similarly, HA stabilization occurred during the adaptation of pH1N1 to humans (35, 36). HA stabilization has been shown to be necessary for airborne transmission of pH1N1 and swine H1N1 gamma viruses in ferrets, in part by increasing longevity of expelled virions (37–39).

The polymerases from avian IAVs are relatively inefficient at genome replication in mammalian cells (40, 41). Therefore, they require adaptive mutations in their polymerase complexes to replicate and transmit in mammals (42–44). Mammalian-adaptive PB2 mutations include T271A, G590S, Q591R, E627K, and D701N (5). PB2-E627K, the best-characterized mutation, was present in the 1918 H1N1, 1957 H2N2, and 1968 H3N2 pandemic viruses and has been associated with avian H7N9 and H5N1 influenza virus adaptation in humans and other mammalian hosts (45, 46). The 2009 pH1N1 virus lacked the PB2-E627K mutation but had phenotypically similar mutations PB2-G590S/Q591R, which were located near residue 627. The swine H1N1 gamma viruses in the present study have mammalian-preferred PB2-G590S/Q591R mutations and avian-like PB2-E627 and PB2-D701.

Influenza pandemic risk assessment algorithms by the US Centers for Disease Control and Prevention (CDC) and World Health Organization (WHO) explicitly consider HA receptor-binding specificity and indirectly account for other human-adaptive viral traits through airborne transmission studies (47, 48). Ferrets are currently considered the gold-standard animal model to study transmission of influenza viruses (20, 49).

Previously, we studied the replication and transmission of gamma-clade H1N1 isolates G15-HA1-mixed and P4-PA-mixed in groups of 3 donor, contact, and airborne ferrets (39). These isolates are A/swine/Illinois/2A-1213-G15/2013 (H1N1) and A/swine/Illinois/2B-0314-P4/2014 (H1N1), respectively. They both contain swine gamma genes in the PB2, PB1, PA, HA, NA, and NS segments and H1N1pdm09 genes in the NP and M segments (39). In the polymerase genes, G15-HA1-mixed and P4-PA-mixed differed at PB2-648 and PA-271; additionally, P4-PA-mixed contained minor variants at PA-321 and PA-386 (Fig. 1A). With respect to HA, P4-PA-mixed contained an HA1-S210 residue (H3 numbering) and had an HA activation pH of 5.5. G15-HA1-mixed contained a mixed population of 85% HA1-N210 and 15% HA1-S210 and had an HA activation pH of 5.8. G15-HA1-mixed had 3/3 airborne transmission events ferrets, while P4-PA-mixed had only 1/3 airborne transmission. As these viruses contained variations and mixed populations in the HA and polymerase proteins and the number of animals per group (n=3) was low, the relative importance of HA stability and polymerase activity in the airborne transmissibility of these viruses was unclear.

**Fig. 1.**
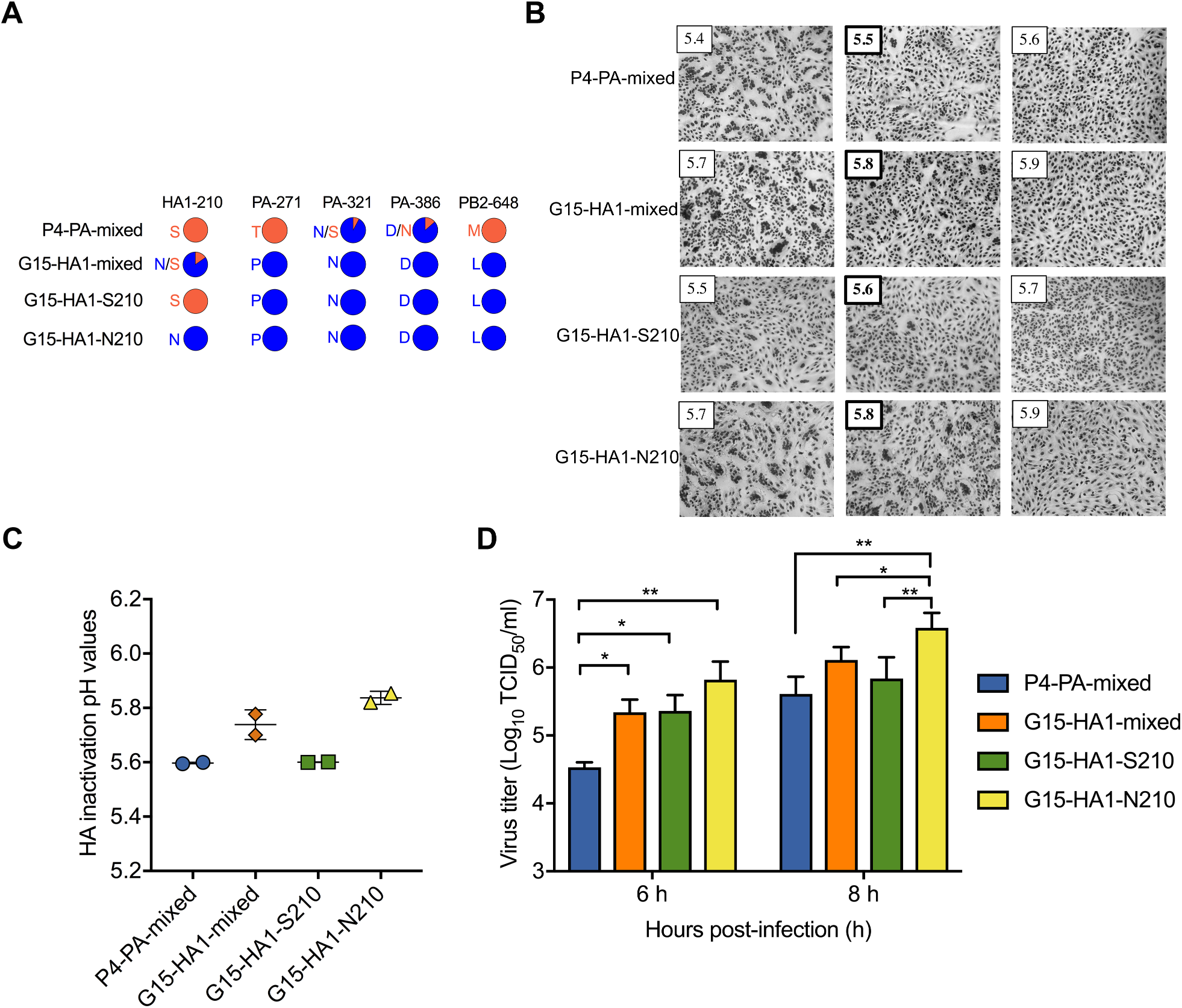
Virus characterization *in vitro*. (A) Sequence variations. Whole genomes of gamma viruses P4-PA-mixed, G15-HA1-mixed, G15-HA1-S210, and G15-HA1-N210 were obtained by next**-**generation sequencing. Amino-acid variations with frequencies ≥ 5% and reads ≥ 10 are shown in colored pie charts. H3 numbering was used for the HA protein. (B) HA activation pH values measured by syncytium assay. Viruses were inoculated into Vero cells at an MOI of 3 PFU/cell. Representative images are shown with the HA activation pH in bold. (C) Virus inactivation pH values. Viruses were treated with pH-adjusted PBS, re-neutralized, and subjected to measurement of residual virus infectivity by TCID_50_. (D) Virus growth in MDCK cells. Viruses were inoculated into MDCK cells at an MOI of 2 PFU/cell. Cell-free culture supernatants were harvested at 6- and 8-hpi and titrated by TCID_50_. All experiments were independently performed at least twice. *P* values were determined according to one-way ANOVA followed by a Tukey’s multiple comparisons test. * and ** represent *P* < 0.05 and 0.01, respectively.

In the present work, we isolated the minor variant G15-HA1-S210 and compared its properties to G15-HA1-mixed and P4-PA-mixed in cultured cells and in ferrets. We also characterized the genotypes and phenotypes of viruses recovered from ferret nasal washes. Overall, enhanced HA stability and polymerase activity were associated with airborne transmissibility, and a stabilized HA appeared to be more stringently required for replication and transmission.

## RESULTS

### Impact of HA and polymerase variations on HA stability and virus replication in MDCK cells

Whole-genome sequencing analyses showed variations at only five amino-acid positions between P4-PA-mixed and G15-HA1-mixed in their entire genomes (Fig. 1A). In the polymerase genes, P4-PA-mixed contained PA-T271, 92% PA-N321 (8% PA-S321), 86% PA-D386 (14% PA-N386), and PB2-M648, while G15-HA1-mixed contained PA-P271, PA-N321, PA-D386, and PB2-L648. In HA, the two isolates differed at HA1-210 with P4-PA-mixed containing HA1-S210 and G15-HA1-mixed containing 85% HA1-N210 (15% HA1-S210). From G15-HA1-mixed, we isolated the majority population (G15-HA1-N210) and the minor variant (G15-HA1-S210). Residue HA1-210 interacts with residues across the trimer interface in the head domain, so the HA1-S210 variation most likely stabilizes the HA protein by resisting dissociation of the heads (39).

We measured the pH of HA activation of the four viruses by syncytia formation assay (Fig. 1B). Vero cells infected with G15-HA1-S210 were triggered to cause syncytia at a highest value of pH 5.6, while G15-HA1-N210 was activated at pH 5.8. Thus, HA1-S210 was more stable than HA1-N210. P4-PA-mixed contained HA1-S210 and induced syncytia at pH 5.5. As 85% of G15-HA1-mixed contained the less-stable HA1-N210 variation, infection with this isolate resulted in syncytia formation at pH 5.8.

The pH of virus inactivation was measured by exposing virus aliquots to media of varying pH, neutralizing, and measuring residual infectivity. G15-HA1-S210 and P4-PA-mixed (which contained HA1-S210) had midpoints of inactivation of pH 5.6 (Fig. 1C), similar to their activation pH values. G15-HA1-N210 had an inactivation pH of 5.8, the same as its activation pH. G15-HA1-mixed had an inactivation pH of 5.75; therefore, this mix of unstable (85%) and stable (15%) components had just a small shift in overall inactivation pH due to the presence of the minority, more-stable HA1-S210.

Previous reports showed differences in polymerase activity and/or HA stability may affect virus one-step growth (50–52). To explore this, we inoculated viruses into MDCK cells at an MOI of 2 PFU/cell and measured the virus titers of the supernatants at 6- and 8-hours post-infection (hpi). Compared to P4-PA-mixed, the three other viruses had higher titers at 6 hpi (Fig. 1D). As P4-PA-mixed and G15-HA1-S210 have identical HA proteins, enhanced replication at 6 hpi was mapped to differences in the polymerase genes. At 8 hpi, G15-HA-N210 had higher titers than the other three viruses. G15-HA1-N210 and G15-HA1-S210 were identical except for HA1 residue 210. Therefore, the HA1-N210 variation (HA activation pH 5.8) enhanced replication at 8 hpi in MDCK cells compared to HA1-S210 (pH 5.6). Overall, both the HA variation and the polymerase variations were found to modulate the production of infectious virus in MDCK cells at relatively early time points when inoculated at an MOI of 2 PFU/cell.

### G15-HA1-S210 had higher polymerase activity than P4-PA-mixed

Virus polymerase activity can be assessed by the accumulation of viral mRNA, cRNA, and vRNA, which are produced by virus transcription and replication (53). To study the impact of the PB2 and PA variations on polymerase activity, we compared G15-HA1-S210, P4-PA-mixed, and P4-PA-purified. G15-HA1-S210 and P4-PA-mixed had identical HA genes, differed at PA-271 and PB2-648, and P4-PA-mixed also had minor variants at PA-321 and PA-386 (Fig. 1A). To exclude the effect of P4-PA-mixed minor variants at PA-321 and PA-386, we obtained purified P4 virus without the above minor mixed populations. Thus, P4-PA-purified and G15-HA1-S210 differed only at PA-271 and PB2-648. The three viruses were inoculated into MDCK cells at an MOI of 2 PFU/cell, and the resulting cell lysates and supernatants were harvested at 4 and 6 hpi. P4-PA-mixed and P4-PA-purified yielded similar levels of mRNA, cRNA, vRNA, and viral titers (Fig. 2); therefore, the minor populations in P4-PA-mixed did not alter virus transcription and replication. Compared to P4-PA-mixed and P4-PA-purified, G15-HA1-S210 had higher mRNA, cRNA, and vRNA levels at 4 and 6 hpi (Fig. 2A-B) and higher viral loads in cell supernatants at 6 hpi (Fig. 2C). In summary, the presence of PA-P271 and/or PB2-L648 in G15-HA1-S210 was associated with higher polymerase activity. These two variations were also present in G15-HA1-mixed and G15-HA1-N210.

**Fig. 2.**
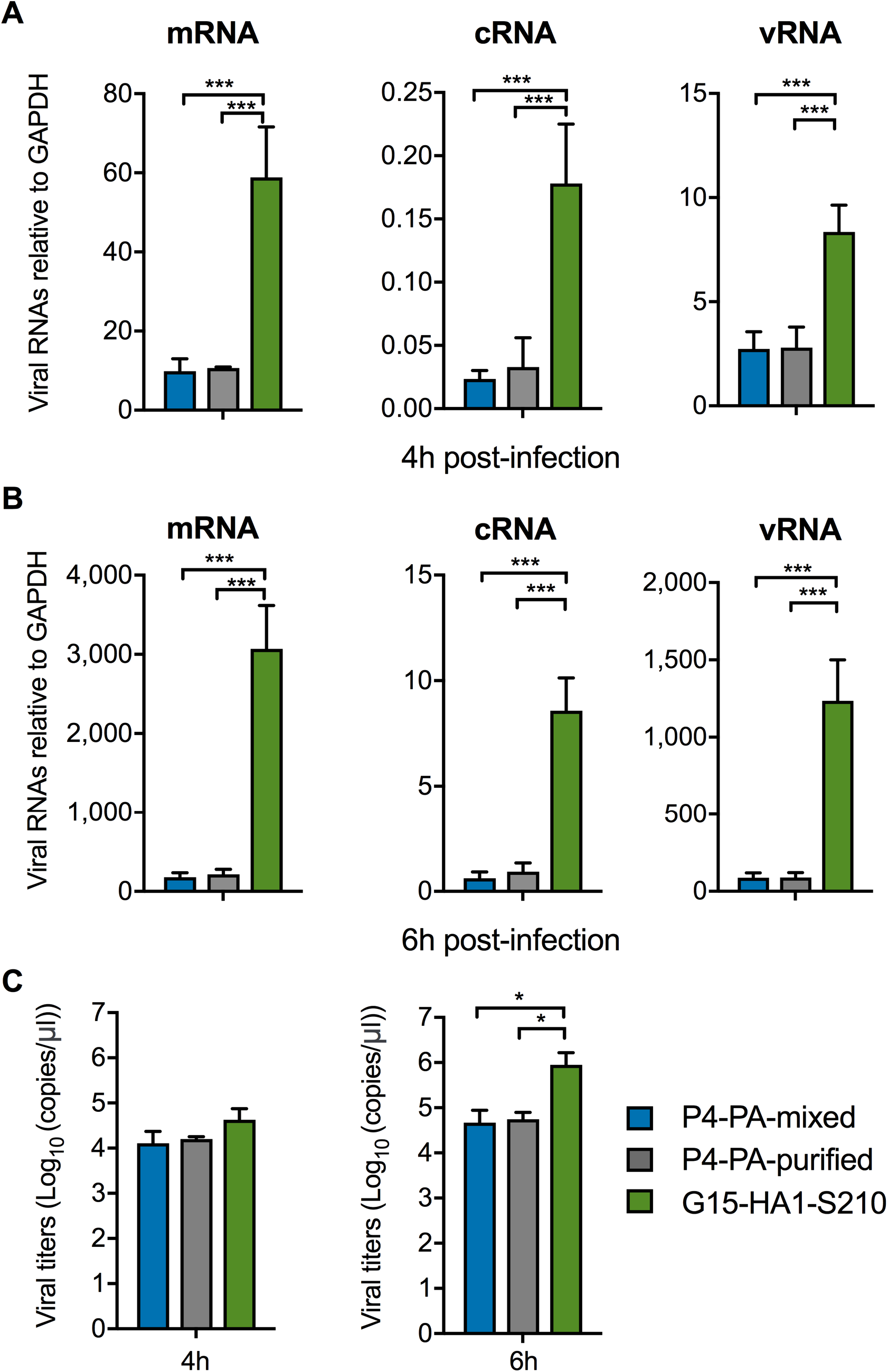
*In vitro* transcription and replication of P4-PA-mixed, P4-PA-purified, and G15-HA1-S210. (A) Virus mRNA, cRNA, and vRNA accumulation after 4 hpi. (B) Virus mRNA, cRNA, and vRNA accumulation after 6 hpi. (C) Virus copies released into culture media. Viruses were inoculated into MDCK cells at an MOI of 2 PFU/cell. At 4 and 6 hpi, the infected MDCK cells were lysed and harvested for total viral RNA. Virus specific mRNA, cRNA, and vRNA were quantified by two-step real-time reverse-transcription PCR using specific primers and normalized to GAPDH mRNA levels. Meanwhile, culture supernatants were harvested for quantification of virions released. Virions released from the cells were quantified as virus copies by real-time quantitative PCR using plasmid PHW2000-NP (G15-HA1-mixed) as a standard. *P* values were determined according to one-way ANOVA followed by a Tukey’s multiple comparisons test. * and *** represent *P* < 0.05 and 0.001, respectively.

### Infection of donor ferrets inoculated with P4-PA-mixed, G15-HA1-mixed, and G15-HA1-S210

G15-HA1-S210 had relatively high polymerase activity and a stabilized HA protein. Compared to G15-HA1-S210, P4-PA-mixed had decreased polymerase activity (due to the presence of PA-P271 and/or PB2-L648) and G15-HA1-mixed had higher HA activation pH and virus inactivation pH (due to 85% HA1-N210) (Fig. 3A). We previously studied the infection of P4-PA-mixed and G15-HA1-mixed in ferrets, using a total of 3 donor, 3 contact, and 3 airborne ferrets for each virus (39). Here, we repeated the experiment and report the combined data for a total of 6 donor, 6 contact, and 6 airborne ferrets, in part to comply with the USDA’s policy on appropriate use of ferrets. The present data is referred to as cubicle 1 and the previous data as cubicle 2. In the present work, we also included two cubicles of ferrets infected with the newly purified virus G15-HA1-S210, thus the ferrets in this group totaled 6 donor, 6 contact, and 6 airborne.

**Fig. 3.**
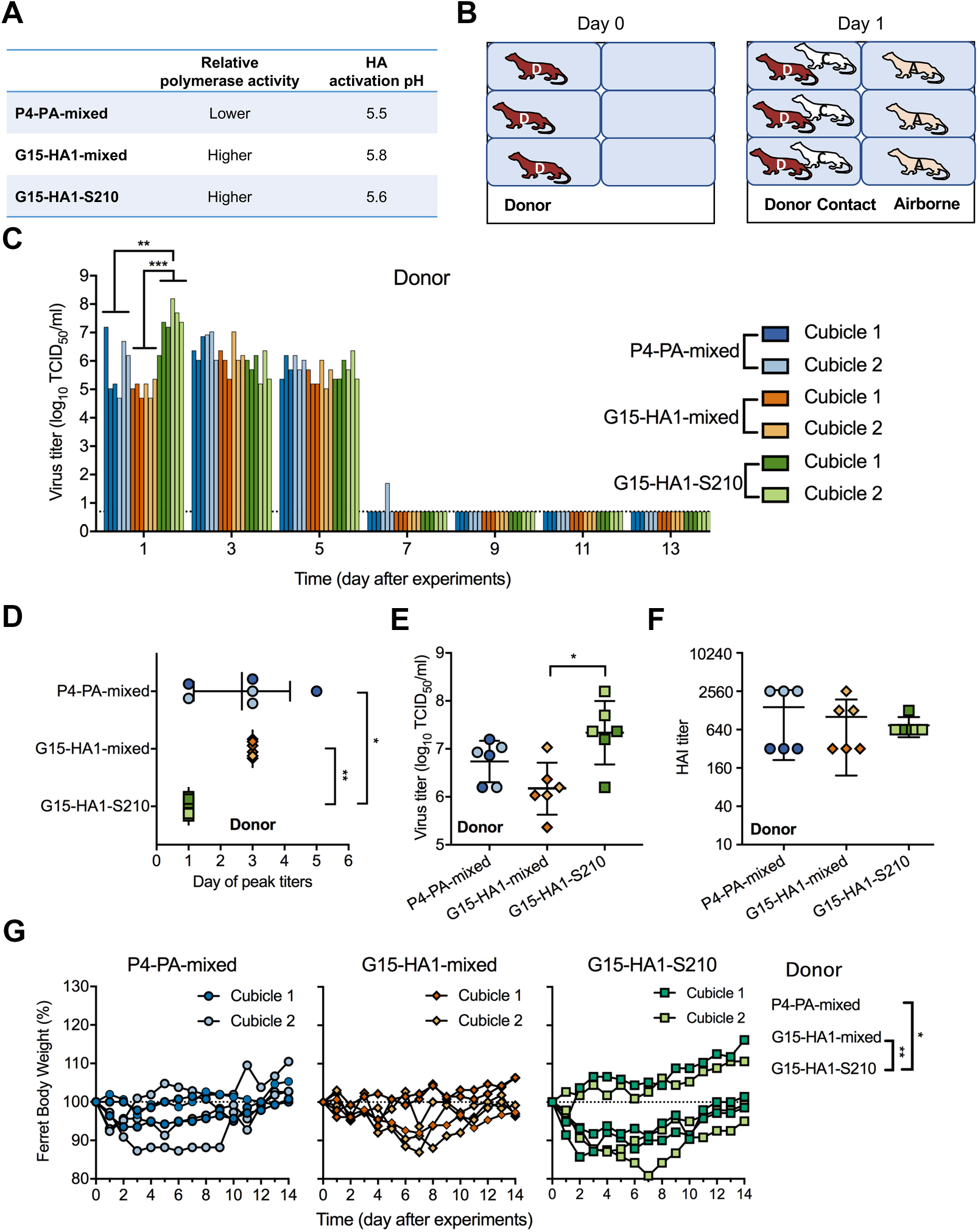
Infection of P4-PA-mixed, G15-HA1-mixed, and G15-HA1-S210 in inoculated donor ferrets. (A) Summary of viral phenotypes. Relative polymerase activity and HA stability for viruses are shown. (B) Ferret experimental caging in each of two cubicles. Viruses were intranasally inoculated into 3 donor ferrets per group on day 0. After one day, 3 naïve ferrets were introduced into the same cages and 3 into the adjacent cages. These ferrets were designated contact and airborne ferrets, respectively. Overall, there were 9 ferrets for each cubicle and two cubicles per virus. (C) TCID_50_ titers of virus in nasal washes from donor ferrets. Ferret nasal washes were collected every other day until day 14. Each bar represents the virus titer of a sample from an individual ferret. For each virus, the first three bars correspond to cubicle one and the next three bars correspond to cubicle two. The bottom dashed lines represent the limit of detection. (D) Day of virus peak titer in donor ferrets. Each symbol represents the day that the virus peak titer was observed in an individual donor ferret. (E) Virus peak titers in donor ferrets. Each symbol represents the virus peak titer in an individual donor ferret. (F) HAI titers of donor ferret sera. (G) Percent body weight of donor ferrets. Ferret body weight was monitored daily until day 14. *P* values were calculated using Mann-Whitney U test. * and ** represent *P* < 0.05 and 0.01, respectively.

In each cubicle, three ferrets were directly inoculated with 10^6^ PFU of viruses, and one day later we introduced three naïve contact and three naïve airborne ferrets (Fig. 3B). Each virus group was evaluated in an isolated cubicle and personal protective equipment was changed before moving between cubicles to avoid cross contamination of viruses. Each cubicle was set to a temperature of approximately 22 °C and relative humidity of 40-60%. All ferrets were monitored for body weight and temperature daily, and ferret nasal washes were collected every other day until day 14. Ferret sera were collected on day 21.

On day 1, G15-HA1-S210 donor ferrets had an average nasal wash titer of 2.18 ×10^7^ TCID_50_/ml, a value more than 10-fold higher than that of P4-PA-mixed (6.88 ×10^5^ TCID_50_/ml, *P* = 0.0065) and 100-fold higher than G15-HA1-mixed (1.08 ×10^5^ TCID_50_/ml, *P* < 0.0001) (Fig. 3C and Table 1). G15-HA1-S210 had greater virus growth after 1 day infection in ferrets than G15-HA1-mixed, while virus replication was similar for the two viruses after 6-8h infection in MDCK cells. This was consistent with previous results showing that HA stabilizing mutations enhanced replication by resisting extracellular inactivation in the mildly acidic upper respiratory tract, whereas MDCK extracellular media were buffered at neutral pH (6, 34). G15-HA1-S210 had peak titers on day 1, while G15-HA1-mixed and P4-PA-mixed had mean peak titers on day 3 and 2.7, respectively (Fig. 3D). Additionally, the average peak titer of G15-HA1-S210 in donor ferrets (2.18 ×10^7^ TCID_50_/ml) was significantly higher than that of G15-HA1-mixed (1.48 ×10^6^ TCID_50_/ml, *P* < 0.5) (Fig. 3E). Overall, G15-HA1-S210, which had a stabilized HA and enhanced polymerase activity, had earlier and higher growth in the nasal cavities of inoculated donor ferrets.

**Table 1.** Virus characteristics before and after infection and transmission in ferrets.

| Virus characterization |  | P4-PA-mixed | G15-HA1-mixed | G15-HA1-S210 | G15-HA1-N210 |
| --- | --- | --- | --- | --- | --- |
| **Genotype variations** | HA1-210 | S | 85% N, 15% S | S | N |
|  | PB2-648 | M | L | L | L |
|  | PA-271 | T | P | P | P |
|  | PA-321 | 92% N, 8% S | N | N | N |
|  | PA-386 | 86% D, 14% N | D | D | D |
| **Phenotype variations in vitro** | HA activation pH <sup>a</sup> | 5.5 | 5.8 | 5.6 | 5.8 |
|  | HA inactivation pH <sup>b</sup> | 5.6 | 5.75 | 5.6 | 5.8 |
|  | Polymerase activity | lower | higher | higher | higher |
|  | One-step growth | lower | higher | higher | highest |
| **Donor ferrets** | Virus growth (log<sub>10</sub> TCID<sub>50</sub>) | 5.8 (± 1) | 5.0 (± 0.28) | 7.3 (± 0.67) | - <sup>f</sup> |
|  | Seroconversion | 3/3; 3/3 <sup>d</sup> | 3/3; 3/3 | 3/3; 3/3 | - |
|  | Day of peak titers | 2.7 (± 1.5) | 3 | 1 | - |
|  | Peak titers (log<sub>10</sub> TCID<sub>50</sub>) | 6.7 (± 0.43) | 6.2 (± 0.54) | 7.3 (± 0.67) | - |
|  | Major variants | PA-S321 | HA1-S210 | none | - |
|  | HA activation pH range | 5.5-5.6 | 5.5-5.9 | 5.5-5.6 | - |
| **Contact ferrets** | Seroconversion | 3/3; 3/3 | 3/3; 3/3 | 3/3; 3/3 | - |
|  | HA activation pH range | 5.5-5.6 | 5.6-5.9 | 5.5-5.6 | - |
|  | Major variants | PA-S321 | HA1-S210 | none | - |
| **Airborne ferrets** | Days for virus shedding | 8-14; 10-14 | -; 8-12 | 8-14; 2-6 | - |
|  | Seroconversion | 3/3; 1/3 | 2/3; 3/3 | 3/3; 1/3 | - |
|  | Major variants | PA-S321 | HA1-S210 | sporadic | - |
|  | Polymerase activity | higher & lower | higher | higher | - |
|  | HA activation pH <sup>e</sup> | 5.5-5.6 | ~5.6 | 5.5-5.6 | - |

<sup>a</sup> HA activation pH measured by syncytia assay.
<sup>b</sup> HA inactivation pH measured by treating viruses with indicated pH and titrating resulting virions by TCID<sub>50</sub>.
<sup>c</sup> The Number of isolates is listed for those grown successfully in MDCK cells after recovery from ferrets in cubicle one and cubicle two.
<sup>d</sup> Data for each virus is listed for each cubicle with the first X/Y number representing cubicle one and the second X/Y representing cubicle two.
<sup>e</sup> HA activation pH of airborne-transmitted virus on first day of isolation.
<sup>f</sup> -, Not measured.

All ferrets seroconverted without significant differences between groups (Fig. 3F). No significant differences in body temperature were detected between the donor ferret groups. Three of six ferrets in the G15-HA1-S210 group had >10% body weight loss for at least two days, compared to only one of six ferrets inoculated with P4-PA-mixed and G15-HA1-mixed (*P =* 0.04 and 0.0055, respectively, Fig. 3G).

### Contact and airborne transmission by P4-PA-mixed

P4-PA-mixed had 6/6 contact transmission and 4/6 airborne transmission as assessed by nasal viral loads and HAI assay (Fig. 4A-B). Airborne transmission events were 3/3 in cubicle one and 1/3 in cubicle two (Table 1). To determine if variants emerged during the experiment, we used next-generation sequencing to determine the whole genomes of viruses from ferret nasal washes. Single-nucleotide variants (SNVs) resulting in amino acid changes were observed for each sample (Fig. 4C). Minor variant PA-S321 represented approximately 8% of the inoculum (Fig. 4D). In cubicle one, PA-S321 was measured at frequencies greater than 10% in all three donors, one contact, and two airborne ferrets (Fig. 4C-D). In airborne ferret A1, PA-S321 was 100% abundant when the virus was first isolated on day 8 and remained at 100% on day 10. On day 8, ferret A1 also had over a dozen minor variants (abundance 5-30%) in HA2, NP, NS1, and NEP. These variants and PA-S321 were also present in airborne ferret A2 on the first day of isolation (day 10), suggesting transmission to A2 may have originated from A1. In both ferret A1 and A2, the minor variants disappeared in the following days. In ferret A2, PA-S321 was approximately 50% abundant on day 10 but was not detectable on day 14. Other variants in PB2, PB1, HA2, NP, NS1, and NEP were either found only in a single ferret or were observed at low percentages. In cubicle two, the frequency of PA-S321 was reduced to an undetectable level in donor and contact ferrets by days 5 and 6, respectively, and PA-S321 was not detected after the single airborne transmission event, which occurred later (by day 12).

**Fig. 4.**
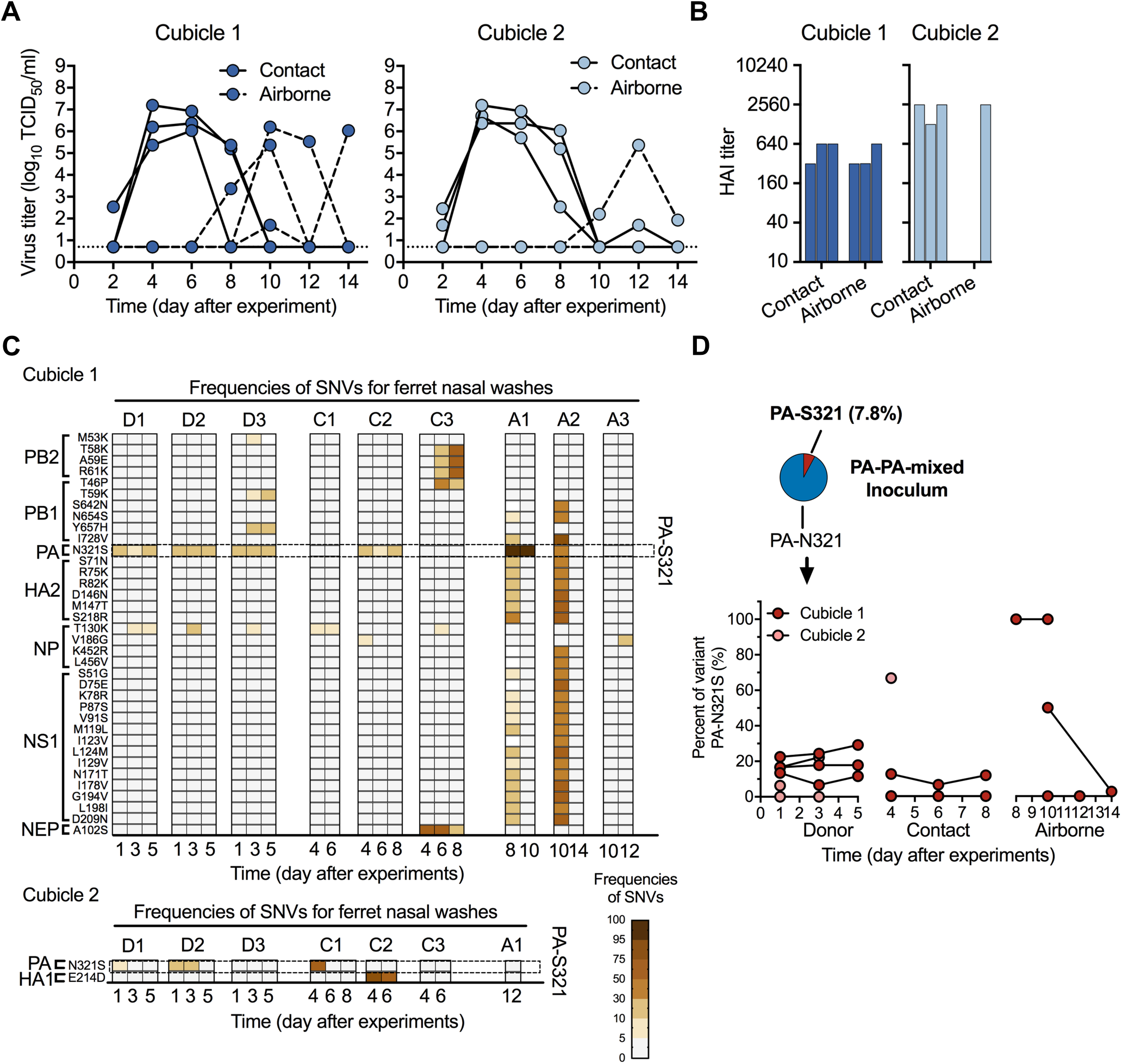
P4-PA-mixed transmission, seroconversion, and frequency of SNVs for nasal washes collected from ferrets after transmission. The ferret experiment was carried out as described in Fig. 3 in which all of the corresponding donor ferrets shed detectable viruses and seroconverted. (A) Virus titers (TCID_50_) of nasal washes from contact (solid lines) and airborne ferrets (dashed lines). (B) HAI titers of day-21 sera from contact and airborne ferrets. (C) Frequencies of SNVs for nasal washes. The whole-genomes of nasal washes collected from contact and airborne ferrets were obtained by next-generation sequencing. SNVs with frequencies ≥ 5% and reads ≥ 10 were read out. The listed SNVs met the following criteria: (1) resulted in protein sequence changes, and (2) appeared in nasal washes in each cubicle ≥ 2 times and/or frequencies greater than 30%. The frequencies of SNVs are represented by the colored boxes (scale bar at bottom right). Each column represents the SNVs for one ferret nasal wash. (D) The proportions of PA-S321 in P4-PA-mixed from the original inoculum and infected/exposed ferrets.

### PA-S321 enhanced polymerase activity

PA-N321S has been identified as an important human-adaptive mutation for avian H5 influenza A viruses (54, 55). Therefore, to assess the effect of the PA-S321 variation on polymerase activity, we plaque purified P4-PA-S321. Compared to P4-PA-mixed, which contained approximately 92% PA-N321 and 8% PA-S321, P4-PA-S321 had higher levels of mRNA, cRNA, vRNA, and viral titers at 4 and 6 hpi in MDCK cells (Fig. 5). This showed that the PA-S321 variant had enhanced polymerase activity. P4-PA-S321 lacked PA-P271 and/or PB2-L648, which were previously shown to enhance the polymerase activity of G15-HA1-S210 (Fig. 2). However, P4-PA-S321 had similar mRNA, cRNA, vRNA, and viral titers at 4 and 6 hpi in MDCK cells as G15-HA1-S210 (Fig. 5), showing that the PA-S321 variation compensated for the lack of PA-P271 and/or PB2-L648.

**Fig. 5.**
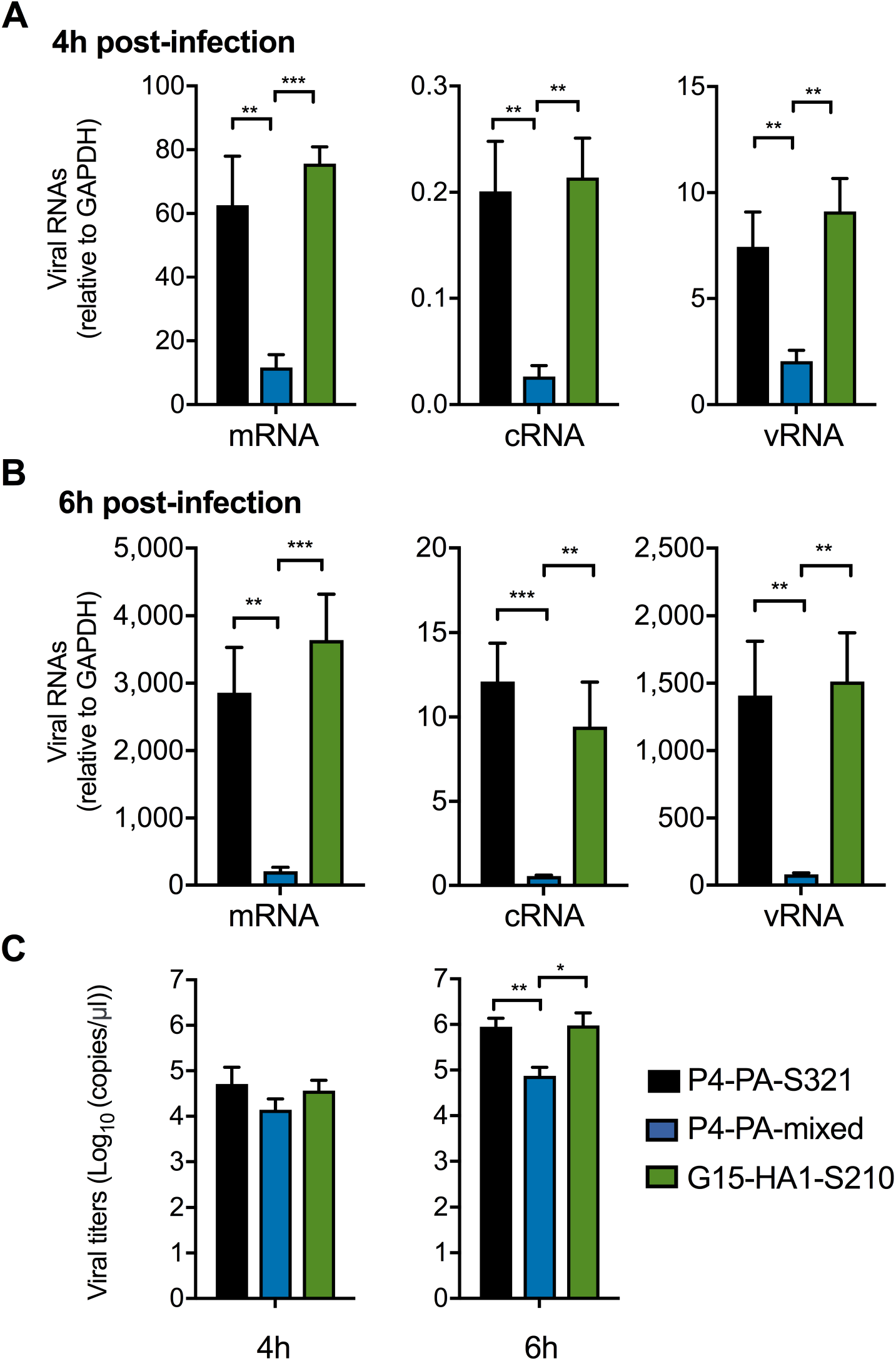
Minor variant PA-S321 had enhanced polymerase activity. (A-C) Evaluation of polymerase activity of P4-PA-S321. The experiment was performed as described in Fig. 2. The transcription and replication of virus containing 100% variant PA-S321 were compared to those of G15-HA1-S210 and P4-PA-mixed. The experiments were independently performed twice. *P* values were determined according to one-way ANOVA followed by a Tukey’s multiple comparisons test. *, **, and *** represent *P* < 0.05, 0.01, and 0.001, respectively.

### Contact and airborne transmission of G15-HA1-mixed and G15-HA1-S210

For G15- HA1-mixed, contact transmission occurred in 6/6 ferrets (100% in both cubicles), and airborne transmission occurred in 5/6 ferrets (2/3 in cubicle one and 3/3 in cubicle two) as detected by seroconversion (Fig. 6B). However, virus was only recovered from 2 airborne ferrets (both in cubicle two) (Fig. 6A). The inoculum of G15-HA1-mixed contained approximately 15% HA1-S210, yet HA1-S210 became enriched to a level greater than 90% in all donor ferrets within three days and was transmitted to all six contact animals (Fig. 6 C-D). We were only able to perform next-generation sequencing on the nasal washes from one airborne ferret (A1), and the virus isolated from this ferret contained 100% G15-HA1-S210, the stabilizing variant.

**Fig. 6.**
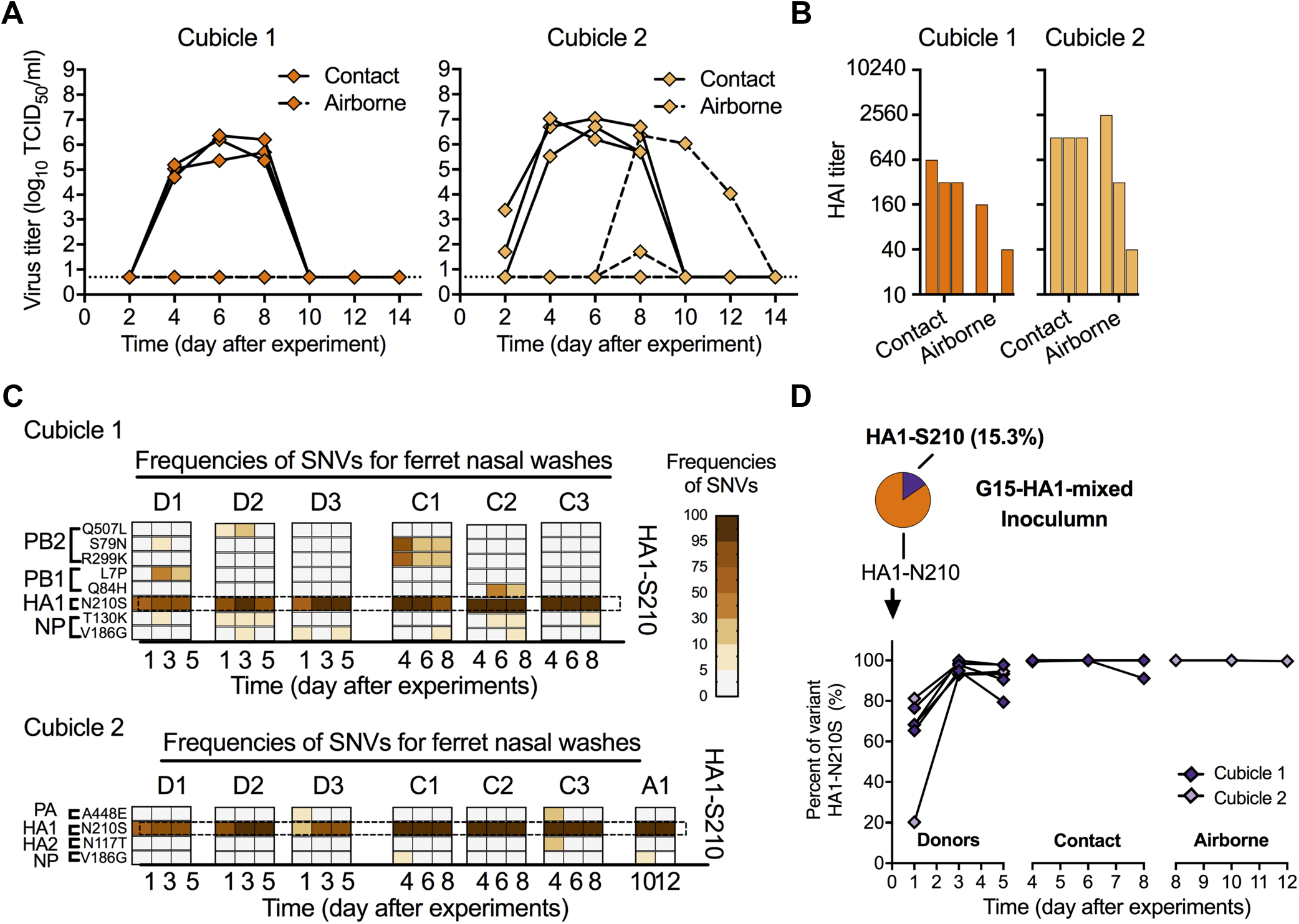
G15-HA1-mixed transmission, seroconversion, and frequency of SNVs for nasal washes collected from ferrets after transmission. The ferret experiment was carried out as described in Fig. 3 in which all of the corresponding donor ferrets shed detectable viruses and seroconverted. The frequency of SNVs were determined and listed as described in Fig. 5C. (A) Virus titers (TCID_50_) of nasal washes from contact and airborne ferrets. (B) HAI titers of day-21 sera of contact and airborne ferrets. (C) Frequencies of SNVs for nasal washes. (D) The proportions of HA1-S210 in G15-HA1-mixed from the original inoculum and infected/exposed ferrets.

Transmission studies were next performed with the G15-HA1-S210 virus. For G15-HA1-S210, contact transmission occurred in 6/6 ferrets (100% in both cubicles), and airborne transmission occurred in 4/6 ferrets (3/3 in cubicle one and 1/3 in cubicle two) (Fig. 7A-B). The HA1-S210 variation remained at 100% in all virus samples sequenced. Virus was recovered from one airborne ferret in cubicle two on days 2-6 and three airborne ferrets in cubicle one between days 8-14 (Fig. 7A). The G15-HA1-S210 group contained a number of minor variants in donor, contact, and airborne ferrets (Fig. 7C). PB1-E751STOP was the most abundant variant isolated from airborne ferrets but was only detected in 2/4, in one of those was not present upon first detectable infection (ferret A1, day 8, cubicle one), and was not detected in nasal washes from the donor and contact ferrets (Fig. 7C). Therefore, this variant was not likely associated with transmission. Ferret A2 in cubicle one contained NP-V352M, NP-E443G, and NA-D311N variations, although these were not detected in the nasal washes of the other airborne viruses or those from donor and contact animals. There were no significant differences in body weight and temperature among the contact and airborne groups.

**Fig. 7.**
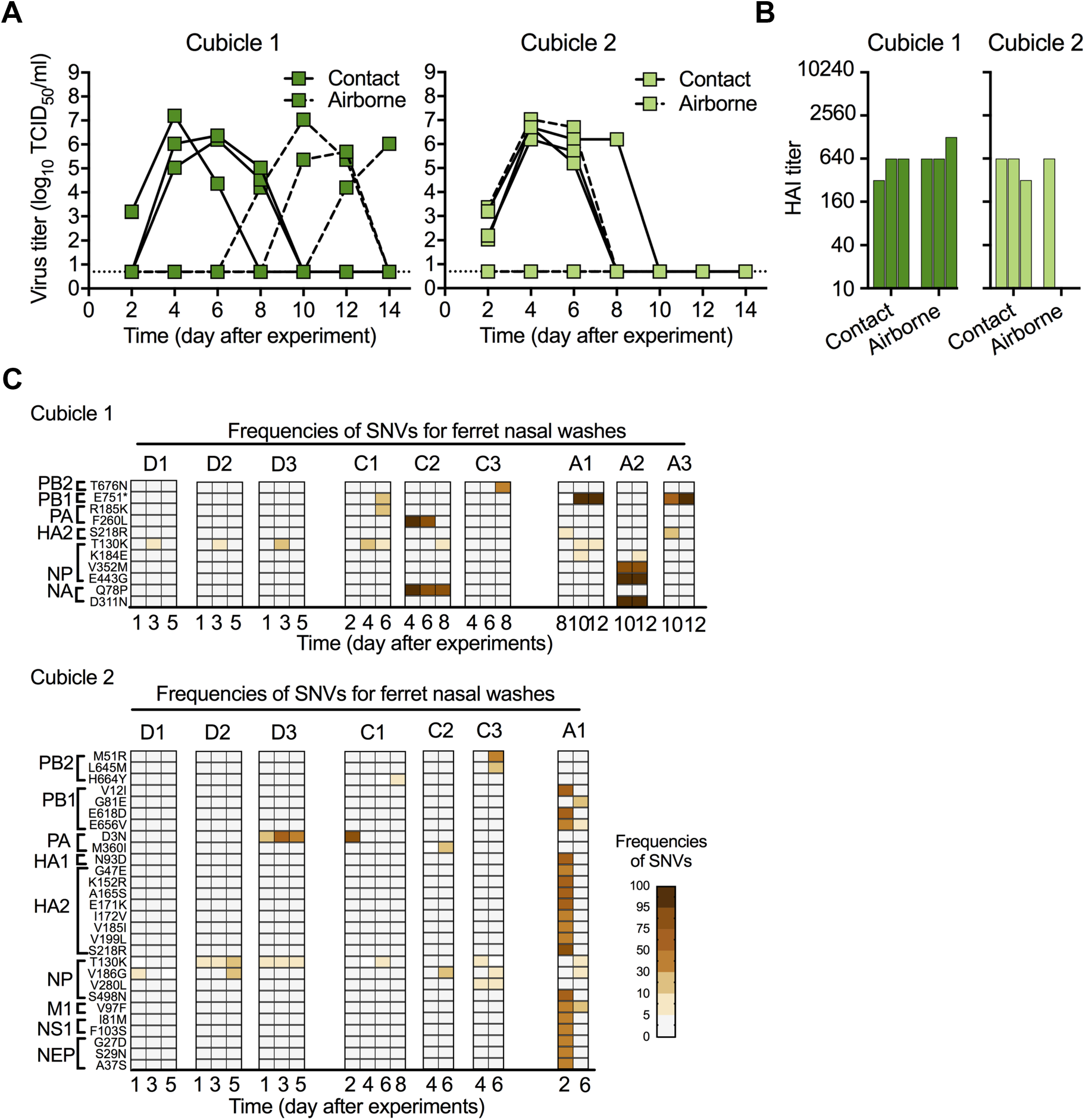
G15-HA1-S210 transmission, seroconversion, and frequency of SNVs for nasal washes collected from transmitted ferrets. The ferret experiment was carried out as described in Fig. 3A, and frequency of SNVs were achieved and listed as described in Fig. 5C. (A) Virus TCID_50_ titers of nasal washes from contact and airborne ferrets. (B) HAI titers of day-21 sera of contact and airborne ferrets. (C) Frequencies of SNVs for nasal washes.

### HA activation pH values of viruses isolated from ferrets

P4-PA-mixed contained 100% HA1-S210, had an inoculum with an activation pH of 5.5, and all recovered virus samples from this group (donor, contact, and airborne) had activation pH values of 5.5-5.6 (Fig. 8A). Similarly, G15-HA1-S210 had a syncytia formation value of pH 5.6, and all recovered virus samples had syncytia formation pH values of 5.5-5.6 (Fig. 8B). Thus, the phenotype of a relatively stable HA protein was maintained for P4-PA-mixed and G15-HA1-S210 after transmission in ferrets. On the other hand, the G15-HA-mixed inoculum contained approximately 85% HA1-N210 and 15% HA1-S210, and the relatively unstable majority component (HA1-N210) contributed to syncytia formation occurring at the relatively high pH of 5.8 (Fig. 1B). Virus isolates obtained from the G15-HA1-mixed donor ferrets, which had mixed populations of HA1-N210 and HA1-S210, had syncytia formation pH values ranging from pH 5.55 to 5.85 (Fig. 8C). Virus isolates from contact ferrets in the G15-HA1-mixed group contained nearly 100% HA1-S210 (Fig. 6D). The three contact ferrets from cubicle one yielded nasal wash samples with HA activation pH values of approximately 5.6-5.7 (Fig. 8C). The single contact ferret from which virus samples were obtained in cubicle two had higher HA activation pH values despite containing HA1-S210 (Fig. 8C). This was most likely due to the presence of a minor population of HA2-N117T, which was previously shown to destabilize the HA protein (39). Airborne-transmitted viruses recovered from G15-HA1-mixed ferret A1 had an HA activation pH of approximately 5.6 (Fig. 8C).

**Fig. 8.**
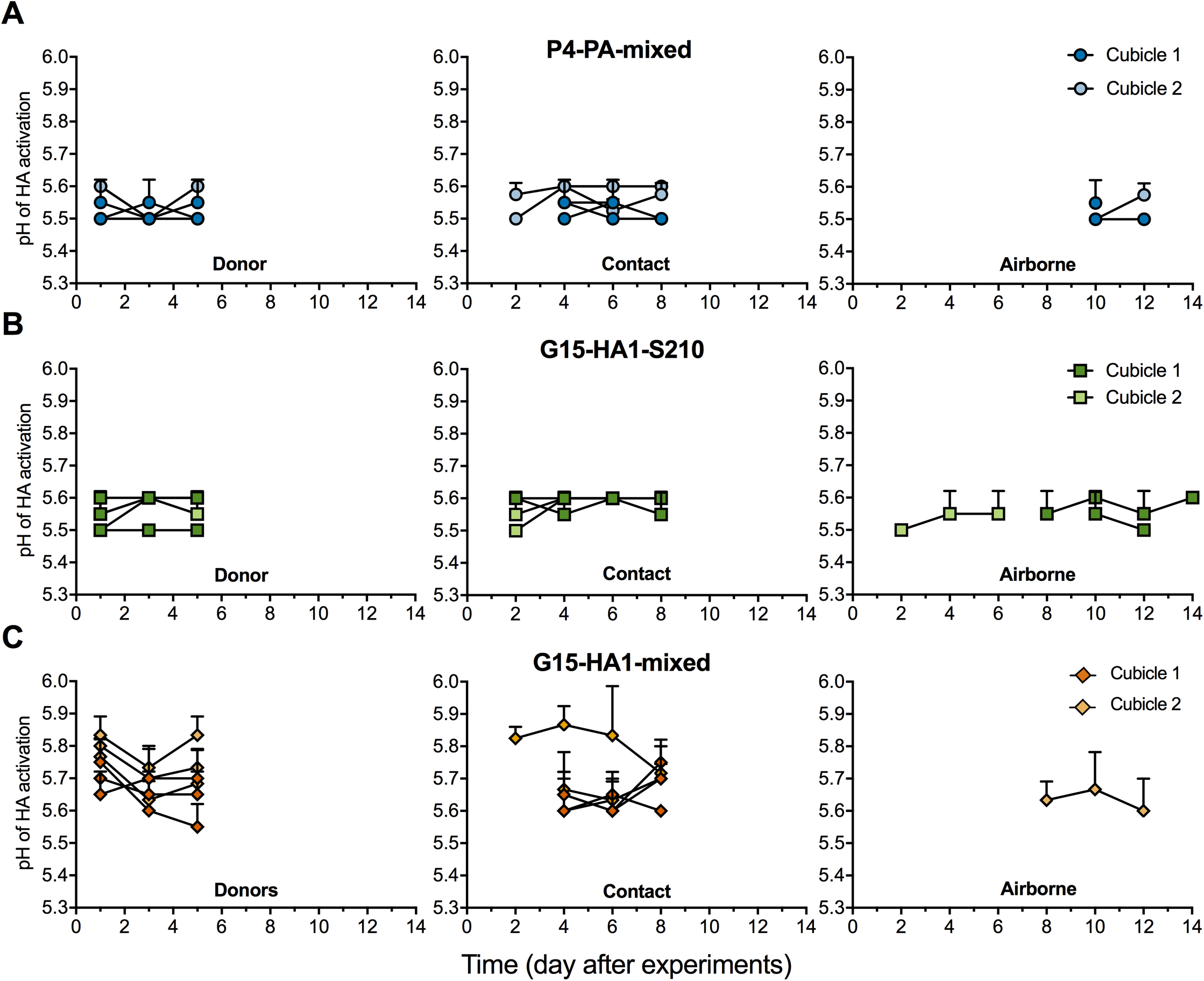
HA activation pH values of viruses isolated from ferrets. (A-C) HA activation pH values of virus-containing isolates from P4-PA-mixed (A), G15-HA1-S210 (B), and G15-HA1-mixed (C). Virus isolates from nasal washes were propagated in MDCK cells once. Syncytium assays were performed to measure the HA activation pH values.

In summary, the G15-HA1-mixed inoculum contained 15% HA1-S210 and this HA-stabilized variant was selected within inoculated ferrets within days and was then transmitted by contact and airborne routes. For all groups studied, all detectable airborne-transmitted viruses had HA activation pH values of approximately 5.5-5.6 and none retained an HA activation pH of 5.8. The P4-PA-mixed inoculum contained 8% PA-S321. This polymerase-enhancing variant remained as a minor population in donor and contact ferrets. PA-S321 was detected in 2/4 airborne-transmitted ferrets. Overall, a stable HA and an efficient polymerase were associated with airborne transmission, and a stabilized HA protein was more stringently selected in ferrets than enhanced polymerase activity.

## DISCUSSION

Three viral traits thought to be necessary for human-to-human and ferret-to-ferret transmissibility by influenza A viruses are HA receptor-binding specificity for *α*-2,6-linked sialic-acid receptors, an HA protein relatively resistant to acid-induced activation/inactivation, and a polymerase complex capable of efficient replication in mammalian cells (5, 17). The swine H1N1 gamma viruses studied here have been shown to bind *α*-2,6-linked receptors with high affinity in addition to having lower-level binding to *α*-2,3-linked receptors (39). P4-PA-mixed had a stabilized HA protein but lacked an efficient polymerase, and G15-HA1-mixed had an efficient polymerase but contained a majority of relatively unstable HA proteins. Both virus isolates contained minority variants at a level of 8-15% abundance that overcame the respective deficiency. Polymerase-enhancing variants were observed in nasal washes from half of recoverable airborne-transmitted P4-PA-mixed, and the HA-stabilizing variant in G15-HA1-mixed was rapidly selected in inoculated donor ferrets and transmitted. Compared to P4-PA-mixed and G15-HA1-mixed, variant G15-HA1-S210, which contained both a stabilized HA protein and an efficient polymerase, had increased early growth in the nasal cavities of inoculated ferrets. Overall, the data demonstrated that the combination of HA stability and polymerase efficiency promotes efficient airborne transmissibility of swine H1N1 gamma viruses in ferrets and that minor variants containing both traits can be selected and airborne transmitted when the majority of a virus sample lacks a stabilized HA or an efficient polymerase.

Multiple traits are required for IAV airborne transmissibility in ferrets, and it is important to understand if these traits must be acquired in a particular order or if they may evolve in any order. For IAVs already possessing *α*-2,6 receptor-binding specificity, the present study revealed two evolutionary pathways for the enhancement of ferret airborne transmissibility: (A) from a virus quasispecies that has an efficient polymerase, a minor variant with a stabilized HA protein may be selected, or (B) from a virus quasispecies with a stabilized HA protein, a minor variant with an enhanced polymerase may be selected.

With respect to pathway A, several other studies showed that an IAV containing *α*-2,6 receptor-binding specificity and an efficient polymerase, but having a relatively unstable HA protein, can acquire airborne transmissibility in ferrets by acquiring an HA-stabilizing mutation. Wild-type 2009 pH1N1 has an HA activation pH of 5.5, and a destabilizing HA1-Y17H mutation increased this value to 6.0 (37, 52, 56). The HA1-Y17H mutant was attenuated in ferrets and did not airborne transmit until acquiring stabilizing HA2-R106K and HA1-H17Y mutations, which decreased the HA activation pH to 5.3 (37). Mechanistically, HA stabilization was shown to increase replication in human airway cells (57), dampen Type I interferon responses in dendritic cells (52), and increase the survivability of infectious virions after respiratory expulsion in ferrets (38). In another study, a recombinant virus was generated that contained the HA from A/Vietnam/1203/2004 (H5N1) with two HA mutations that conferred α2,6 receptor-binding specificity, and the other seven genes from A/California/04/2009 (H1N1) with a PB2-E627K mutation that enhanced polymerase activity (27). This virus and an adaptive mutant containing an HA1-N158D mutation that deleted a glycosylation on the HA head were incapable of airborne transmissibility in ferrets. Instead, a final mutation needed for the acquisition of airborne transmissibility was HA1-T318I in the stalk, which decreased the HA activation pH to 5.3. When HA-stabilizing mutations were introduced into other H5N1 viruses lacking α2,6 receptor-binding specificity, virus growth in the nasal cavities of ferrets was enhanced but airborne transmission did not occur (29, 30, 58).

With respect to pathway B, P4-PA-mixed had a stabilized HA protein and its airborne transmission in 2/4 cases was associated, at least initially, with the presence of the PA-S321 variation that enhanced polymerase activity. A/Indonesia/5/2005 (H5N1) acquired airborne transmissibility in ferrets by a combination of mutagenesis and adaptation that enhanced receptor-binding activity (HA-T156A and HA-Q222L or HA-G224S), polymerase activity (PB2-E627K and PB1-H99Y), and HA stability (HA-H103Y) (28, 59). These mutations arose at similar steps during adaptation, and viruses lacking any of the single mutations HA-T156A, PB2-E627K, PB1-H99Y, or HA-H103Y, or both the HA-Q222L or HA-G224S mutations, were loss-of-function for airborne transmission in ferrets. In the case of reassortant IAVs containing genes from the human 1918 virus and avian H1N1 viruses, the human HA gene enabled efficient contact transmission but did not allow airborne transmission in ferrets (20). Instead, the human 1918 PB2 protein was found to be both necessary and sufficient to enable ferret airborne transmission in the context of a virus that contained the human 1918 HA and NA genes. Thus, while selection of enhanced polymerase activity in the present work was less stringent than selection of a stabilized HA, the adaptation of IAVs for airborne transmission in ferrets, and presumably in humans, has been associated with both HA stability and polymerase activity and both evolutionary pathways (A and B) appear viable.

Quasispecies have been associated with IAV fitness and virulence in various hosts (60, 61). In this study, minor variants of swine H1N1 viruses with enhanced polymerase activity or HA stability were selected in ferrets and associated with airborne transmissibility. The transmission of minor variants has been previously observed for IAVs in humans (62), equines (63), and guinea pigs (64). This suggests minor variants, in addition to the majority population of a virus, need to be considered when assessing the risk of IAVs for human pandemic potential. Next-generation sequencing has been used to track the outgrowth of HA-stabilizing variants in ferrets in this study and others (37–39, 56). Deep sequencing of samples from H7N9-infected patients and their surrounding poultry/environment showed acquisition of the polymerase-enhancing PB2-E627K variation in humans (45). H3N2 and H1N1 growth in mice has been shown to not necessarily proceed through linear accumulation of adaptative mutations (65). Overall, as next-generation sequencing becomes cheaper, faster, and more efficient, real-time monitoring of human-adaptive variations may enable better preparedness for future outbreaks.

Numerous studies have been performed to identify and characterize human- and ferret-adaptive mutations in IAVs. Examples include those focusing on the polymerase complex (42, 51, 66–72) and the HA protein (38, 56, 58, 73–81). In the present study, we found G15 isolates containing PB2-L648/PA-P271 had higher polymerase activities than P4-PA-mixed, which contained PB2-M648/PA-T271. These mutations may lead to increased production of viral pathogen-associated molecular patterns (PAMPs) or defective interfering particles as well (82–84). PB2-L648 was previously shown to increase polymerase activity of an H10N8 virus (85). The present study also showed that PA-S321 increased polymerase activity. PA-S321 has previously been identified as an adaptative marker for H5N1 influenza A viruses in humans (54, 55, 86). The HA1-S210 mutation was shown to decrease the pH of HA activation and the pH of virus inactivation here and in our previous work (39).

In summary, a combination of efficient polymerase activity and a stabilized HA were found to enhance airborne transmission of swine H1N1 gamma viruses in ferrets, and these two properties may be acquired in either order. Human infections of seasonal influenza viruses have been far less frequent since the start of the SARS-CoV-2 pandemic, mask mandates, and social distancing. Reemergence of seasonal influenza has begun to occur. Ultimately, exotic IAVs in other species, especially swine, and the potential for cross-species transmission make it essential to monitor potential pandemic IAV strains and to assess their pandemic risk. Swine H1N1 gamma viruses are one of the most prevalent lineages in swine (87), have caused sporadic human infections in recent years (15, 88), and have been shown here to acquire increased transmissibility in ferrets by adaptive HA and polymerase mutations. A recent study has identified a swine Eurasian avian-like influenza virus with pandemic traits (89). Surveillance and risk assessment studies are needed for IAV pandemic preparedness in addition to studies investigating the evolutionary pathways by which pandemic potential may arise and genetic and phenotypic traits associated therewith.

## MATERIALS AND METHODS

### Cells and viruses

Madin–Darby canine kidney (MDCK) cells were maintained in minimum essential medium (MEM, Thermo Fisher Scientific). African green monkey kidney (Vero) cells were maintained in Dulbecco’s Modified Eagle Medium (DMEM, Life Technologies™). Both culture media were supplemented with 10% HyClone^®^ standard fetal bovine serum (FBS, Life Technologies) and 1% penicillin/streptomycin (P/S, Thermo Fisher Scientific). The cells were grown at 37 °C with 5% CO_2_ (50, 51).

A/swine/Illinois/2A-1213-G15/2013 (H1N1) (G15-HA1-mixed) and A/swine/Illinois/2B-0314-P4/2014 (H1N1) (P4-PA-mixed) were originally isolated from nasal swabs from pigs in commercial swine herds in the United States that showed symptoms of influenza-like illness, as described previously (39, 90). G15-HA1-S210 and G15-HA1-N210 were plaque purified twice from MDCK cells using G15-HA1-mixed. The whole-genome sequences of G15-HA1-S210 and G15-HA1-N210 were verified by next-generation sequencing. P4-PA-purified was plaque purified from P4-PA-mixed and verified by next-generation sequencing to not contain minor mixed populations at PA-321 and PA-386. All viruses were propagated in MDCK cells with 1 μg/ml tosylsulfonyl phenylalanyl chloromethyl ketone (TPCK)–treated trypsin, as described previously (39, 50).

### Syncytium assay

Viruses were inoculated into Vero cells in 24–well plates at a multiplicity (MOI) of 3 PFU/cell. After maintaining at 37 °C with 5% CO_2_ for 16 hours (h), the infected cells were treated with DMEM supplemented with 5 μg/ml TPCK–treated trypsin for 15 minutes (m) and then pH– adjusted PBS buffers for 15 m. Subsequently, the infected cells were maintained in DMEM supplemented with 5% FBS for 3 h at 37 °C. After that, the infected cells were fixed and stained using a Hema 3™ Fixative and Solutions (Fisher Scientific). Syncytia were recorded using a light microscope. The cutoff values for virus HA activation pH were determined at the highest treated pH for which the stained cells in 24-well plates contained more than two syncytia with at least five nuclei, as described previously (39, 52, 91).

### Virus inactivation pH assay

Viruses were treated with a series of pH-adjusted PBS buffers in a ratio of 1:100 for 1 h at 37 °C. The resulting infectious viruses were titrated by TCID_50_ on MDCK cells (30, 37). After obtaining virus titers as a function of exposure pH, virus inactivation pH values were calculated by using GraphPad Prism version 7 software (Graph-Pad Software, San Diego, CA). The parameter LogEC50 was generated using the model Nonlinear Regression [Equation: log (agonist) vs. response-Variable slope (four parameters)] followed by the least squares fitting method.

### Virus growth assay

Viruses were inoculated into MDCK cells in 24–well plates at an MOI of 2 PFU/cell. After incubation for 1 h at 37 °C, the infected cells were washed by PBS twice and then maintained at 37 °C with MEM culture media (1 ml/well) supplemented with 1 μg/ml TPCK-treated trypsin. At 6- and 8-hpi, cell culture supernatant was collected and titrated by TCID_50_ on MDCK cells (37, 92).

### Virus mRNA, cRNA and vRNA quantification

Production of virus RNAs (mRNA, cRNA, and vRNA) during influenza A virus transcription and replication was quantified using a two-step real time RT-qPCR, modified from the method described previously (51, 68, 93). Briefly, viruses were inoculated into confluent MDCK cells in 24–well plates at an MOI of 2. The infected cells were incubated at 37 °C for 1 h, washed by PBS twice, and cultured with MEM supplemented with 1 ug/ml TPCK-treated trypsin (1 ml/well). At 4 and 6 hpi, culture supernatants and infected cells were harvested separately.

To quantify virus mRNA, cRNA, and vRNA, total RNAs were extracted from the infected cells using RNeasy Mini Kit (Qiagen). The corresponding cDNAs of mRNA, cRNA, and vRNA were generated by reverse-transcription PCR (RT-PCR) using Transcriptor First Strand cDNA Synthesis Kit (Roche). Primers with specific tags for differentiations of virus mRNA, cRNA, and RNA were used (mRNA-RT-PCR primer: CCAGATCGTTCGAGTCGTTTTTTTTTTTTTTTTTCTCATGTTTCT; cRNA-RT-PCR primer: GCTAGCTTCAGCTAGGCATCAGTAGAAACAAGGGTGTTTTTTCTC; vRNA-RT-PCR primer: GGCCGTCATGGTGGCGAATTGGCCACAGGATTAAGGAATATC) Real-time quantitation PCR were performed to quantify these three RNA levels with specific primers targeting virus HA gene (mRNA-real time-forward primer (FP): CCAGTTCATTGGTACTGGTAGTC, mRNA-real time-reverse primer (RP): CCAGATCGTTCGAGTCGT; cRNA-real time-FP: CCAGTTCATTGGTACTGGTAGTC, cRNA-real time-RP: GCTAGCTTCAGCTAGGCATC; vRNA-real time-FP: GGCCGTCATGGTGGCGAAT, vRNA-real time-RP: ACCGTACCACCCATCTATCA;). Internal control GAPDH mRNA was also reverse-transcribed and quantified using oligo (dT)_20_ as RT-PCR primer and the following as real-time PCR primers (FP: ATTCCACCCATGGCAAATTC, RP: CGCTCCTGGAAGATGGTGAT).

To quantify viral copy number released in cell-free supernatant, total RNAs were extracted using QIAamp Viral RNA kits (Qiagen). A two-step quantitative real-time reverse-transcription (RT) PCR was applied to quantify viral copy number. The primers used were listed as the following, RT-PCR-primer: AGCAAAAGCAGG; Real-time-FP (targeting NP gene): CTGCTTGTGTGTATGGGCTT; Real-time-RP (targeting NP gene): TGAGCTGGATTTTCATTTGGT. Plasmid PHW2000-NP (G15-HA1-mixed) was used as a standard for quantitative real-time PCR.

### Ferret transmission experiments

Thirty-six male Fitch ferrets (five- to six-month-old, purchased from Triple F Farms, Sayre, PA) were verified to be serologically negative for currently circulating influenza A and B viruses and the swine H1N1 gamma influenza viruses by HAI assay. At the same time, these ferrets were also shown to be serologically negative for currently circulating SARS-CoV-2 by ELISA. The nasal swabs of these ferrets were verified to be negative before the experiments by performing real-time PCR reactions.

The transmission study was performed as described in previous work (39). Ferret body weight and temperature were monitored daily until day 14. Ferret nasal washes were collected every other day until day 14. Additionally, all ferrets were euthanized on day 21 to collect whole blood. HAI assays were performed to examine seroconversion of ferret sera using the corresponding inoculated virus in each group (37, 39, 91). The presence or absence of the stabilizing mutation HA1-N210S did not alter the results of HAI assays in control experiments. Animal studies were performed in compliance with St. Jude Children’s Research Hospital Animal Care and Use Committee guidelines under protocol 459.

### Virus whole-genome sequencing and analyses

Virus whole genomes were obtained by next-generation sequencing (39, 56, 91). Viral RNAs were extracted from ferret nasal wash samples or cell-free supernatants using QIAamp Viral RNA kits (Qiagen). The corresponding cDNAs were achieved by reverse–transcription PCR using SuperScript III First-strand Synthesis System (Thermo Fisher Scientific). DNA library was prepared by PCR amplification using Phusion High-Fidelity PCR Master Mix with HF Buffer (New England BioLabs). The resulting PCR products were purified by QIAquick Gel Extraction Kit (Qiagen) and submitted to St Jude Harwell Center for virus whole-genome sequencing.

Sequencing data were analyzed using CLC Genomics Workbench version 11.0.1. SNVs resulting amino-acid variations were reported when the predefined quality scores were met and present in forward and reverse reads at an equal ratio. All variants presented in this study were supported by at least 10 reads with a minimum frequency of 5% (unless otherwise stated). Heat maps and parts of whole were generated by using GraphPad Prism version 7 software (GraphPad Software, San Diego, CA). The frequency of a SNV is calculated by the percent of read counts of a nucleotide variant to the whole nucleotide read counts at a specific position.

### Statistical analysis

All data analyses were performed using GraphPad Prism version 7. One-way ANOVA followed by a Tukey’s multiple comparisons test, or Mann-Whitney U test were used to determine statistical significances. *P* value < 0.05 was considered significant.

## ACKNOWLEDGMENTS

We thank Guohua Yang, Ashley Webb, and Melissa Penaflor for help with ferret experiments. We also thank St. Jude Animal Resources Center (ARC) for help with ferret experiments. We thank the Hartwell Center DNA Sequencing & Genotyping, Hartwell Center Functional Genomics, and the Hartwell Center Genome Sequencing Facility. This work was funded, in part, by the National Institute of Allergy and Infectious Diseases under Centers of Excellence for Influenza Research and Surveillance (CEIRS) contract no. HHSN272201400006C, St. Jude Children’s Research Hospital, and the American Lebanese Syrian Associated Charities (ALSAC). The content is solely the responsibility of the authors and does not necessarily represent the official views of the National Institutes of Health.

## CONFLICT OF INTEREST

The authors declare no competing financial interests.

## Notes

### Competing Interest Statement

The authors have declared no competing interest.

